# Successive pattern in winter wheat main stem structure modeling

**DOI:** 10.1101/2024.07.22.604639

**Authors:** Jingru Yin, Chen Zhu, Qing Li, Pengyan Li, Chaoyu Fan, Yangmingrui Gao

## Abstract

With the rise of research on functional structural plant models (FSPMs), it is particularly important to realistically describe the structural traits of a plant. In 3D FSPMs, the true structural information of the canopy can make the model’s estimation of functional traits more accurate. Since canopy is the medium of the crop stands that sense and respond to the external environment, accurate estimation of detailed organ sizes in crop canopy is crucial, and current FSPMs still need to be improved. Here the structure of wheat (*Triticum aestivum*) was measured in detail at the phytomer level. Dataset was built which covers a set of ∼100 cultivars and 11 site-year under various field managements such as sowing density, inter-row, nitrogen fertilizer and sowing date. The relationships of successive phytomer sizes were quantified and the variance was analyzed for each independent experiment, only site was found as a significant impact factor. A conceptual model based on the successive pattern was proposed and validated for detailed final phytomer sizes. Then we coupled the meteorology data - the ratio of average daily gross radiation and temperature during the emergence and architecture mature with the model. The validation has taken place three times to emphasize model’s independent of cultivars, sites dependent and the ability of model parameters deriving from environmental factors, respectively. The results revealed the feasibility of the newly built wheat main stem structure model with the R^2^ higher than 0.85. Compared with ADEL-Wheat and Sirius Quality, the model achieved better performance in blade width and blade area estimation. According to the concise and easily acquired parameters of this model, it has potential for high throughput phenotyping detail wheat canopy structure. Previous work on the relationship between blade width and circumference of apex shoot in maize illustrated that the cell number was the most likely explanation for this pattern, which means the same work holds for similar crops.

## 3. Introduction

Over the past two decades, the functional-structural plant model (FSPM) has been a central focus for plant scientists and well developed from gene to community scale. Deciphering the underlying robust pattern and the root of possible deviation from it should be put high on the agenda in the following years (Louarn and Song 2020). Mimicking the performance of crops has long been considered an efficient way to predict and manage to improve the yield results under various conditions. Thus, Sirius Quality and ADEL-Wheat were established and updated for wheat in-silico experiments (Fournier et al. 2003; Jamieson et al. 1998; Liu et al. 2019; Martre et al. 2015; Martre and Dambreville 2018). However, even the simplified model needs plenty of parameters as input, some of which are not easy to acquire. And the simulation of detailed architecture information like organ size still needs to be optimized and tested. Besides, these models have limited application of cultivars and environments.

Plant morphology is dynamic and interdisciplinary of plant biology. Bucksch et al. (2017) reviewed the challenge of modern morphology study is to quantify the transformations resulting from different underlying genetic, developmental, and environmental cues. And that is based on Goethe’s paradigm-a repetitive process of transformation. Skinner and Nelson (1994) reported the coordination pattern in the grass from the cell division and proved that major transitions in leaf and tiller development appeared to be synchronized among at least three adjacent nodes. There is strong coordination between the main stem and its tillers and between the different phytomers (FSPM elementary units) depending on their positions along the stem (Tivet et al. 2001). Martre and Dambreville (2018) confirmed previous observations, according to which the sizes of leaf laminae and sheaths on the main stem and axillary tillers follow the same developmental patterns when the phytomer number is corrected to account for the difference in the final leaf number of each axis. For shoots architecture modeling, the model based on coordination rules for maize proposed by Zhu et al. (2014) and completed by Vidal and Andrieu (2019) was elaborate and robust on grounds of its self-regulation character. Then Gauthier et al. (2020) innovated the model for grass shoots architecture by combining metabolic regulations with coordination rules. Although this model was validated in wheat, comprehensive and robust models for wheat shoots morphology more concisely are in demand.

This paper proposed the successive pattern in wheat which was inspired by a doctoral thesis of Mariem (2016). The dimensions of blade, sheath, and internode have strong linear relationships between phytomer n and n+1. It is derived from coordination rules mainly focusing on the relationship between any two successive phytomers. The properties of phytomer n+1 are conservatively associated with the properties of phytomer n. Similar shreds of evidence were also shown in maize reported by Andrieu et al. (2006) that the length of leaf 10 highly correlated with leaf 11 and the same relationships for the previous 5 leaves are independent of density and cultivar. However, this pattern has not quantified completely and implemented into any model in wheat so far.

Given that the final length of any organ can be used to simulate its kinetics of extension and tillers follow the same growth and development pattern of the main stem (Fournier et al. 2003), we aimed at establishing parametric functions to define the final dimensions (length and width) of phytomer components (blade, sheath, and internode) for all leaf ranks on the main stem which can be fitted into a wide range of situations. To this end, we 1) quantified the correlation between successive phytomers; 2) modeled the structure of the main stem using coefficients; 3) compared the successive pattern model to Sirius Quality and ADEL-Wheat in this study by using a very large dataset.

## 4. Review

### 4.1. ADEL-Wheat

First proposed by Fournier et al. (2003), ADEL-Wheat presented the 3D architecture of wheat, which is derived from thermal time. The final organ sizes were simulated by the functions of phytomer positions along the axis. In total, the number of parameters required to describe the mature length of all organs of a plant is 10 + Nt. Maximum leaf width was found to be well correlated to either sheath length or blade length (depending on the rank), this allows, with the use of four additional parameters, modeling of the leaf width. The model was established based on two experiments and needs to be modified under various conditions.

Mariem (2016) reduced the input parameters of the ADEL-Wheat model by using a representative dataset covering 28 different winter wheat canopies from winter wheat crops grown in Grignon, France with different sowing dates, cultivars, and nitrogen fertilizer rates, in which statistical models were developed to parameterize the final length and shape of leaves.

Liu et al. (2019) represented the linear relationship between lamina width and lamina length from the first phytomer to the order of maximum length of phytomer (typically the penultimate leaf) in ADEL-Wheat, which adapted from the work of Dornbusch et al. (2011). The author described that the lamina width continues to increase although the lamina length decreases from the penultimate to the flag leaf. For lamina length, E.J.M (2002) also reported that it increases with increasing leaf number from the base, reaching a maximum of one or two leaves before the flag leaf after which the length declines so the flag leaf is somewhat shorter than the longest leaf. However, lamina width increases with leaf position so that the flag leaf is the widest. Sheath length also increases with leaf position, markedly so for the culm leaves. Besides, the final sheath length was derived from lamina width based on their strong linear correlation. This general relationship is also reported by Evers et al. (2005).

### 4.2. Sirius Quality

Sirius Quality is a process-based crop growth model that simulates the phenology and canopy development which has been developed and calibrated for spring and winter bread wheat (Jamieson et al. 1998; Martre et al. 2006; Martre et al. 2015).

In the recent work of Martre and Dambreville (2018), Sirius Quality describes the dynamic of leaf area from the growth of individual leaves and tillers using the same approach of ADEL-Wheat. More specifically, the author proposed a hypothesis that a unique relationship may exist between the lamina area and leaf rank for all the tillers, whatever the sowing date or density. Then the experiments covered two cultivars, locations and densities were used to test this hypothesis, in which the whole growing period was divided into three-phase: before floral initiation, after the flora initiation, and from penultimate leaf to the flag leaf. For the juvenile stage, the lamina area was considered constant, and the empirical data was used for different cultivars. According to leaves after floral initiation, the number varies between 4 to 6, which was used as the intercept of the linear relationship between lamina area and the phytomer rank counted basipetally. Besides, the potential lamina area of the penultimate leaf was used as a parameter for the equation of this linear relationship. For the flag leaf, the only empirical ratio between the penultimate leaf and the flag leaf was used to do the estimation.

## 5. Material and methods

### 5.1. Field experiments

We conducted 10 field experiments, four of which locates in China over 3 sites while the others are in Grignon, France (Table 1). Therein, Jiangsu province, where winter wheat (*Triticum aestivum L.*) was monitored over two growing seasons from 2019 to 2021 over three sites (Xuzhou, Jurong, and Baima, respectively correspond to 34°28′N, 117°30′E; 32°18′N, 119°14′E and 31°62′N, 119°20′E) within two subtypes of climate. A large dataset of winter wheat ranged over 7 growing seasons from Grignon, France (48°51′N, 1°58′E) were used in this study.

**Table 1.**
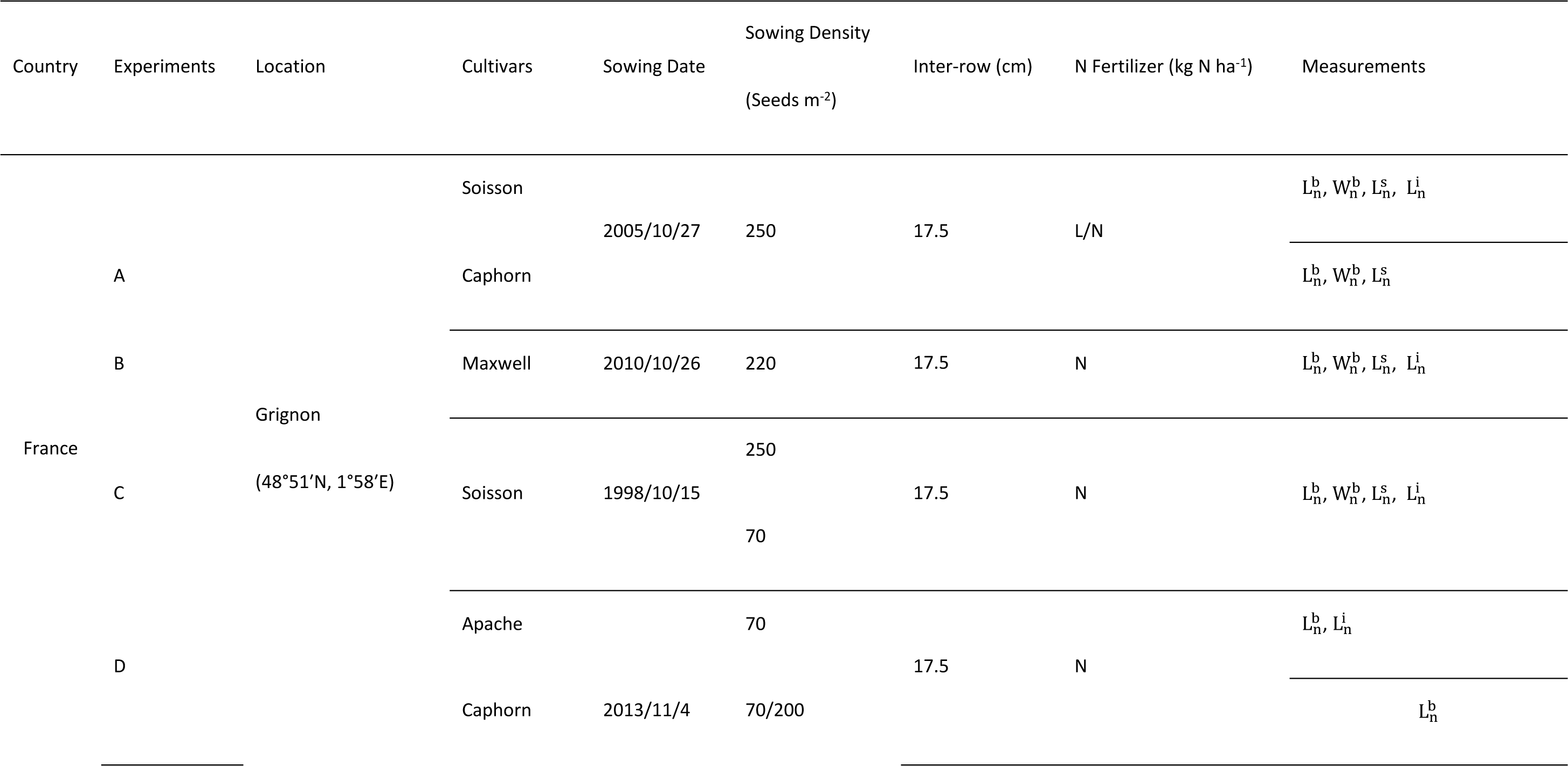
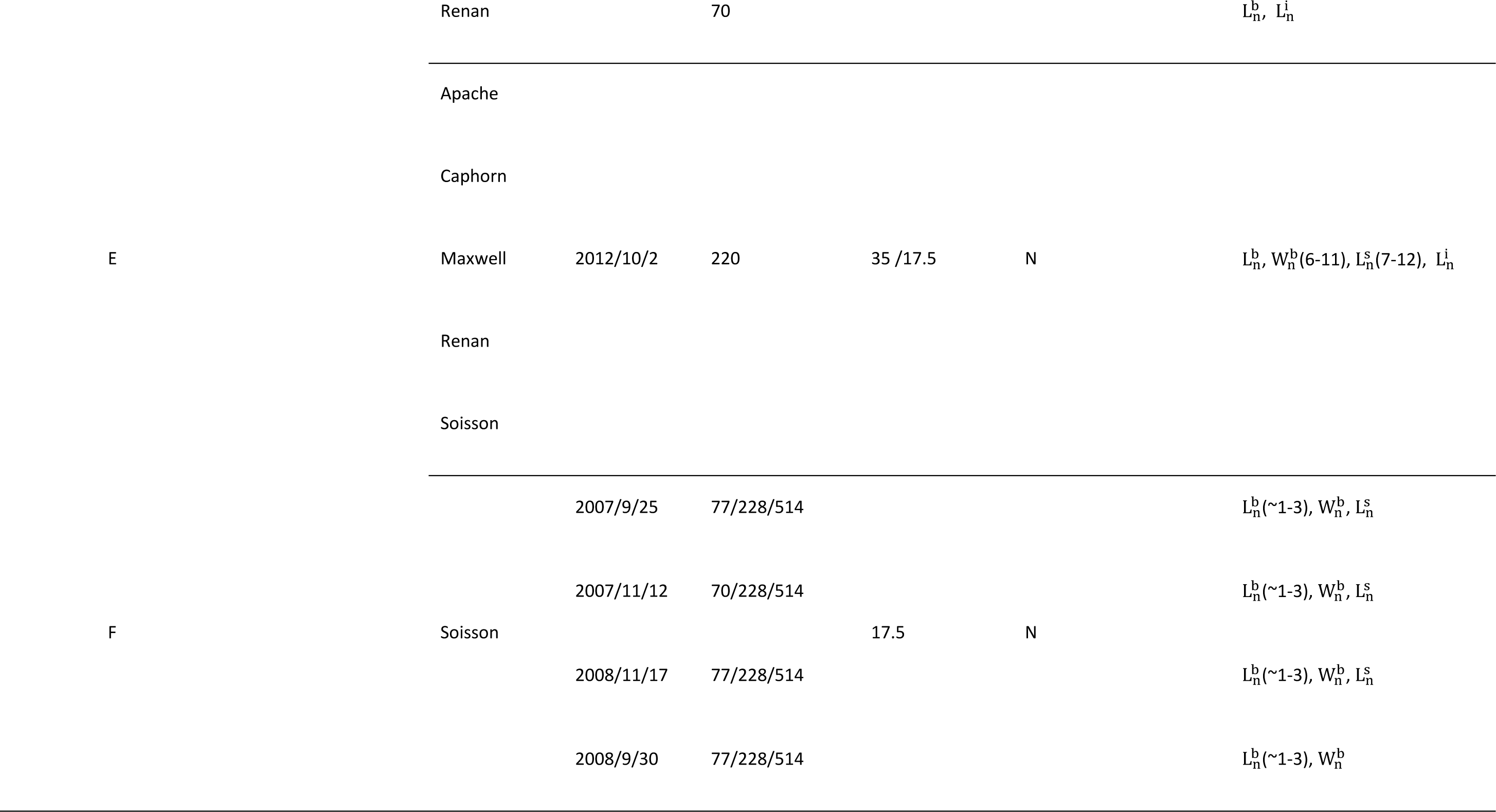
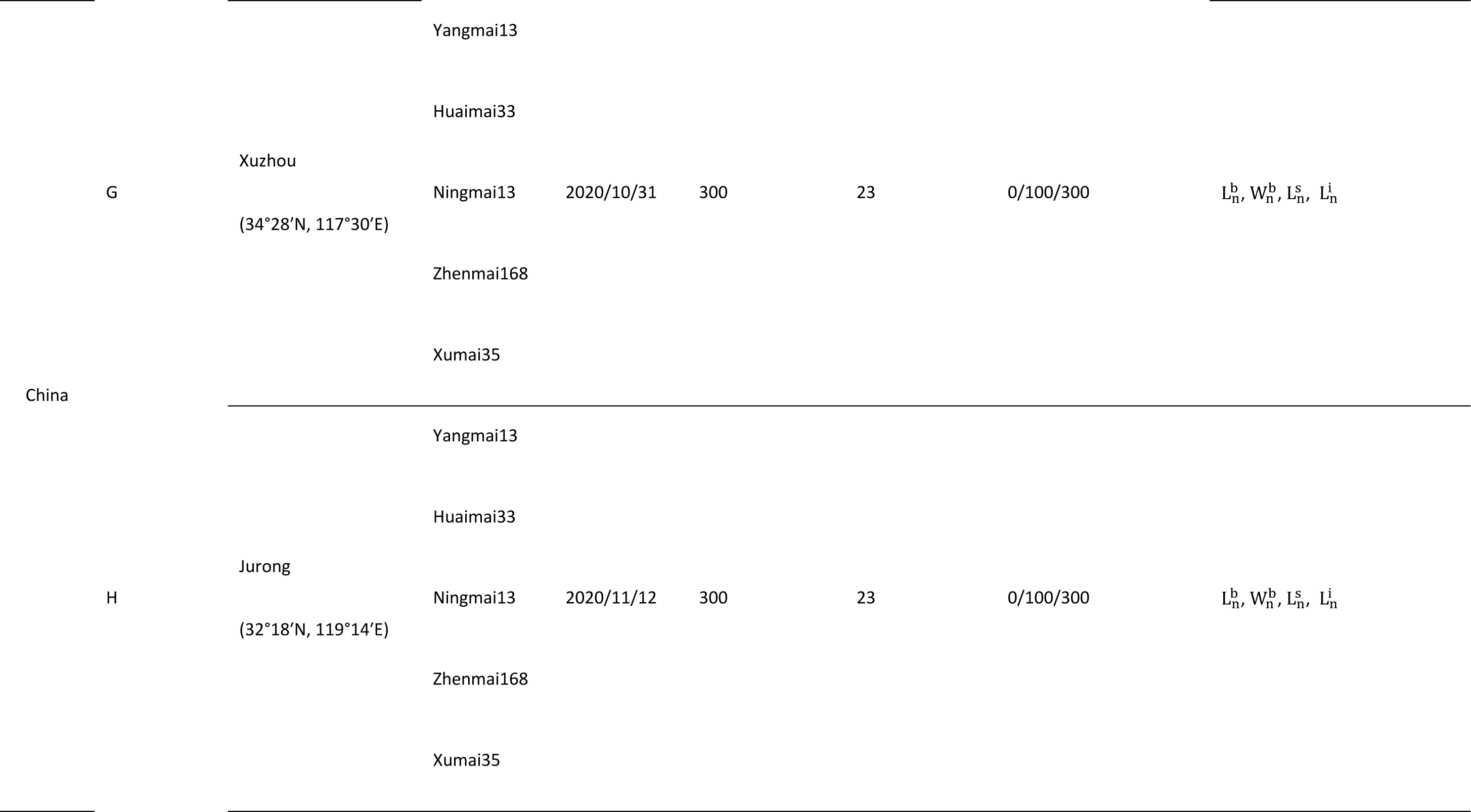
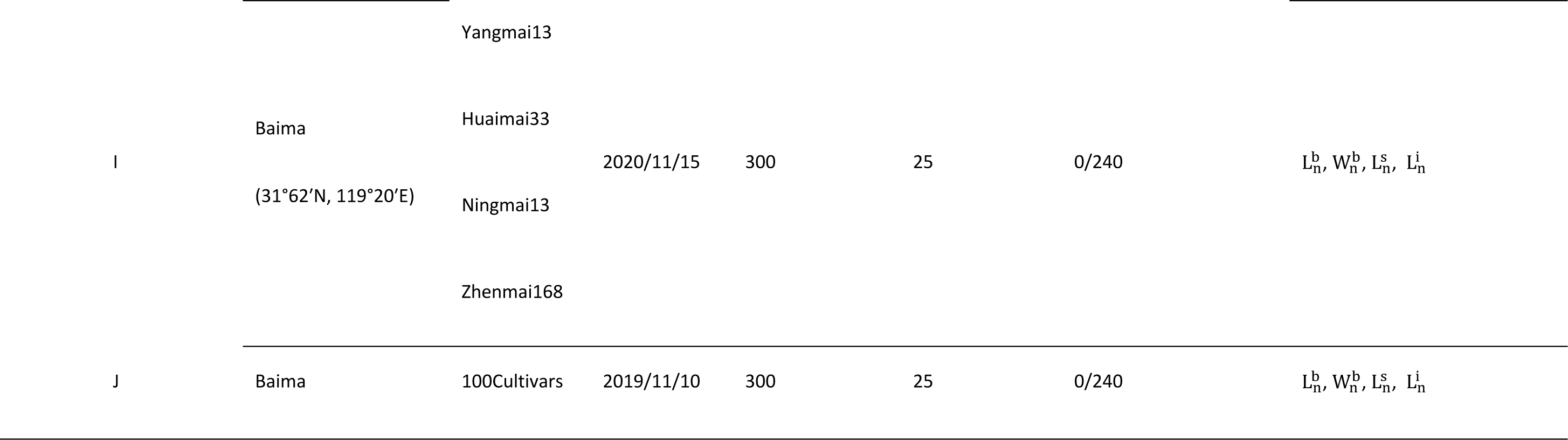
Field experiments and measurements. L: low nitrogen treatments, N: normal nitrogen treatments, 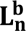: blade length of the phytomer n, 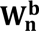: blade ith of the phytomer n, 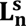: **sheath lenght of the phytomer n,** 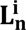: **length of internode n**

Treatments were set differently for 10 independent experiments, which includes 4 locations, 13 sowing dates, around 100 winter wheat cultivars, 2 inter-rows, 3 nitrogen fertilization, 3 sowing densities. A set of ∼100 winter wheat cultivars was sown in mid-November of 2019 in Baima, where the experiment was conducted in a split-plot design under 2 nitrogen treatments with two replications per cultivar using micro plots with a size of 1 m**^2^** and repeated in 2020. Besides, we carried out the same two experiments during the 2020 growing season combined with 5 cultivars and three levels of soil N in the field of Xuzhou and Jurong, in which plots were arranged in a randomized compete for block design of 9 m width by 10 m long. For further information about the French dataset please refer to (Abichou et al. 2018). Plants were grown under non-limiting conditions of water and nutrients and were kept free of disease and weeds by appropriate fungicide and herbicide applications.

### 5.2. Field measurements

We tagged 6∼10 plants per plot, only focusing on the phytomer ranks of the main stem. Several destructive sampling occasions were undertaken during the crop cycle from emergence to anthesis to cover all leaf ranks, in which 10 plants were collected and measured per treatment (Table 1). The final dimensions (width and length) of phytomer components (blade, sheath, and internode) from the main stem were measured manually with a ruler or pasted on paper then scanned into image. Measurements were performed when leaves were fully developed, and no leaves had senesced for each phytomer. Note that due to the quarantine of Covid-19, we measured only 3 tagged plants per plot and missed some phytomers from the rank of 4 to 6 in the 2019 Baima field experiment. Local meteorology data was downloaded from the National Meteorology Information Center.

### 5.3. Data processing and statistical analysis

For each plot, the mean value of the raw data of each phytomer rank was prepared for analysis. And scanned images were processed by the Lamina2Shape software (Dornbusch and Andrieu 2010). The dimensions (width and length) of phytomer components were normalized into 0-1 using the maximum blade length, blade width, sheath length, and internode length of every main stem of wheat respectively. Data processing was performed in software MATLAB R2019b (The Math Works, Inc.). The main fitting algorithm is QR decomposition. For robust fitting, fitlm uses M-estimation to formulate estimating equations and solves them using the method of Iteratively Reweighted Least Squares (IRLS). The mean value of acquired data per treatment (cultivar and management) was used for ANOVA analysis.

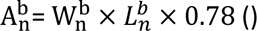

### 5.4. Model Evaluation

For ADEL-Wheat, since the lamina length along the stem is more descriptive other than derived from the other traits based on parameters, only the lamina width and the sheath length were used for model comparison. The formulae are as follows, where 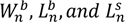 is the blade width, blade length and sheath length of the phytomer *n* respectively. (Adapted from Liu et al. (2019))

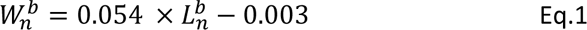

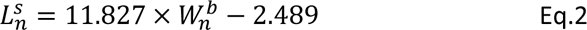

For Sirius Quality, the comparison taken place on the lamina of large leaves (exclude the flag leaf), we adjusted the model using the current dataset based on the following formula. 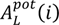 represents potential lamina area of the phytomer *i* (counted from the base),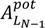 is the potential final lamina area of the penultimate leaf, η is the x-axis intercept of the relationship between 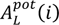 and the phytomer number for large leaves, and *N* is the final leaf number. (Adapted from Martre and Dambreville (2018))

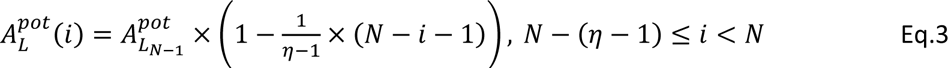

The optimization converged in all cases. Root mean squared errors (RMSE) were calculated and the adjusted R square derived from MatLab Curve Fitting Tool were used to evaluate models in this work.

## 6. Results

### 6.1. Modeling the phytomer dimensions based on the successive phytomers coordination

In this paper, independent dataset of typical cultivar from China and France respectively was selected as the result for representation. The figure shows the upward trend dimension of phytomer compositions along with the development (Fig 1 a-e): On the whole, each factor showed an increasing trend with the phytomer rank, and the representative cultivars in China and France also showed an almost consistent trend. The blade width 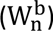, blade length 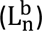, sheath length 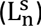, and blade area 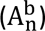 showed a smaller growth rate in the first 4-5 Phytomer ranks. When jointing beginning, internodes appeared, and internode length 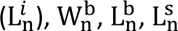 and 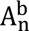 almost simultaneously began to show a significant increasing trend. The difference is that 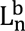 starts to show a downward trend from the penultimate phytomer rank, and at this point, the phytomer rank of these two cultivars from China and France is also the same.

**Fig 1.**
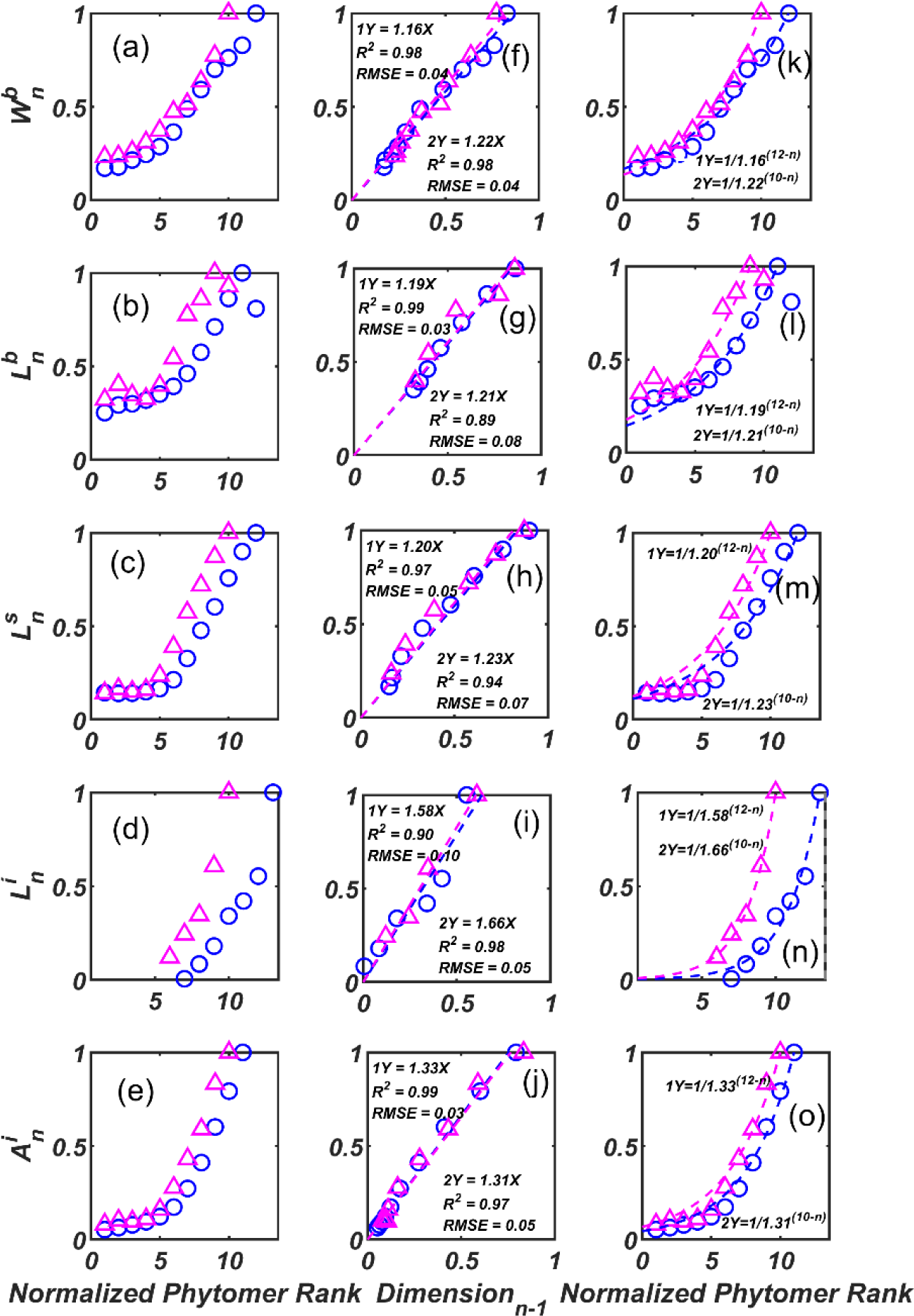
Pattern along the main stem and the coordination of the successive phytomer. (a)-(e) The variance of blade width 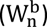, blade length 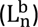, sheath length 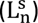, internode length 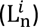, and blade area 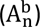 corresponding with normalized phytomer rank. (f)-(j) The correlations between successive phytomer dimensions. (k)-(o) Description of main stem structure based on the successive phytomer dimensions coordination. The pink triangle represents the data from experiment H (Jurong in China, cultivar Ningmai 13 under normal nitrogen treatment), while the blue circle symbols are from experiment C (Grignon in France, cultivar Soisson).

The dimensions 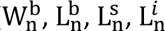, and 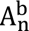) between successive phytomers are highly correlated, and the relationship between them can be well represented by a linear model (Fig 1 f-j, R^2^ > 0.90). The linear coefficient *a* varies in the range of 1.19-1.66, that is, the successive phytomers show positive growth, and coefficient *a* of the two representative cultivars of China and France are almost the same (Δ < 0.1). So, integrate the dimension relationship of successive phytomers to uniformly described by the following formula:

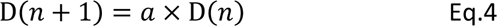

Where *n* is the phytomer rank of the main stem in the range from 1 to N, N is the final phytomer number, D(*n*) is the dimension of *n* th phytomer, and *a* is the slope between dimension *n*+1 and *n*. Detailed coefficient *a* of all treatments showed in the appendix (Table S1). Further, using Eq4 to model the main stem structure, we have:

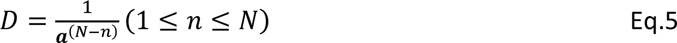

Where *a* represents the parameter, *D* represents the normalized organ dimension, *n* represents the number of phytomer rank with the range of 1 and *N*, and *N* represents the final number of phytomers. Among them, this study models all blade widths and internode lengths, and started from n = 4 for blade length and sheath length, and the results are shown in Fig 1 k-o: Eq.5 can well describe the variance of the successive phytomers dimensions’ changing with the rank. Based on these, we analyzed the impact factors of the successive phytomers coordination law further.

### 6.2. Impact factor of the phytomer dimensions coordination law

We applied Eq.4 to the dataset of all organ dimensions, analyzed and obtained the coefficient *a*, and then performed an analysis of variance (ANOVA) on all independent experiments. ANOVA results of the blade length on *a* indicated that the cultivar and management measures (sowing date, sowing density, nitrogen fertilizer level, row spacing, year) had no significant effect on 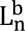, and different sites had significant impacts (Table 2, *P < 0.001*). In addition, the same analysis was performed for other dimensions of the phytomers (leaf width, leaf sheath length, internode length) with similar results (Table S2-4). It was concluded that phytomer dimensions were independent of cultivar and nitrogen treatment, and only meteorology had a significant impact on them. Further, we analyzed the correlation between the dimensions of each phytomer and meteorological factors, and the results showed that blade length and width had higher positive correlations, and the photothermal quotient had a positive correlation with the two (Fig S1, R values were 0.7 and 0.67, respectively).

**Table.2.**
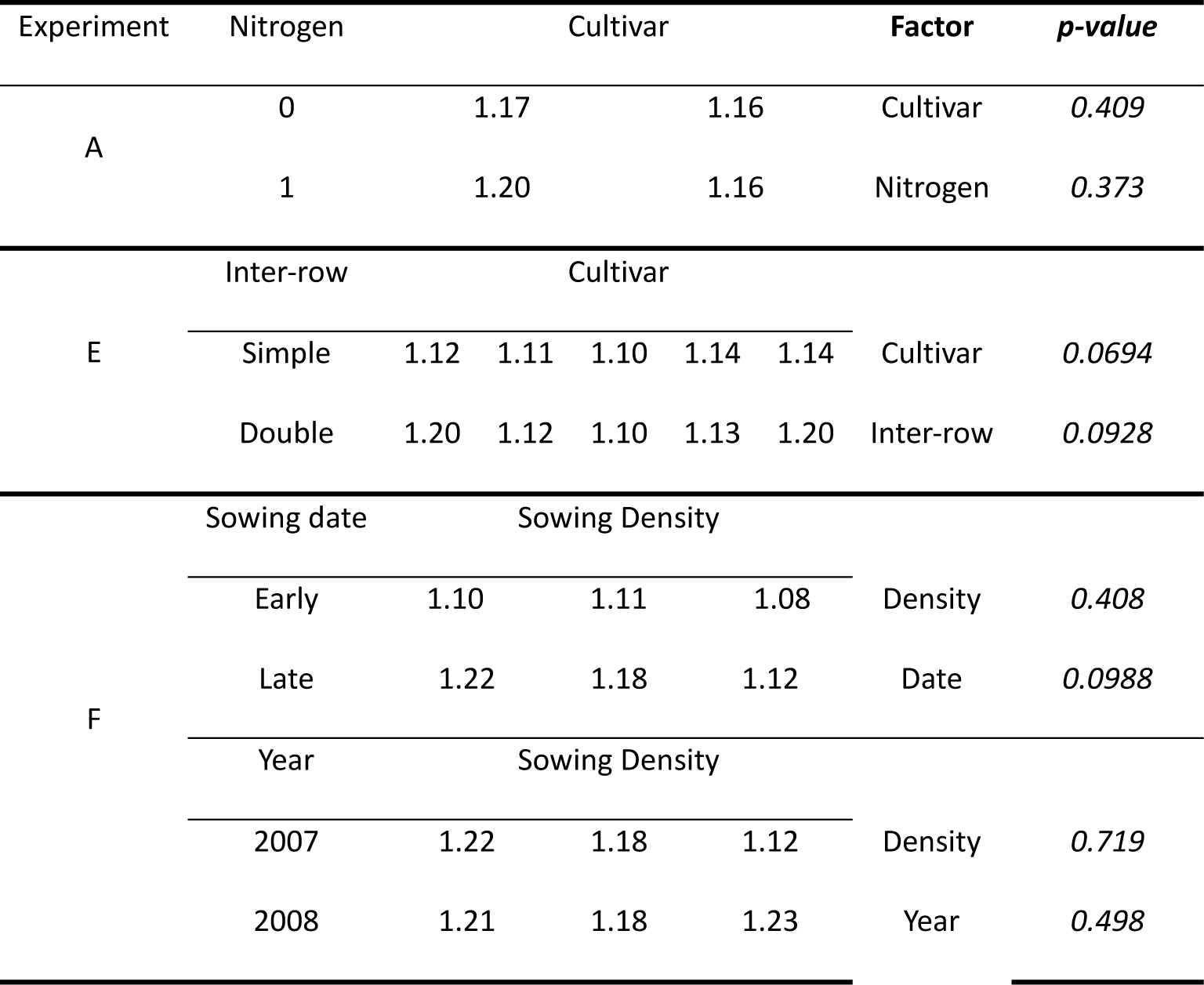
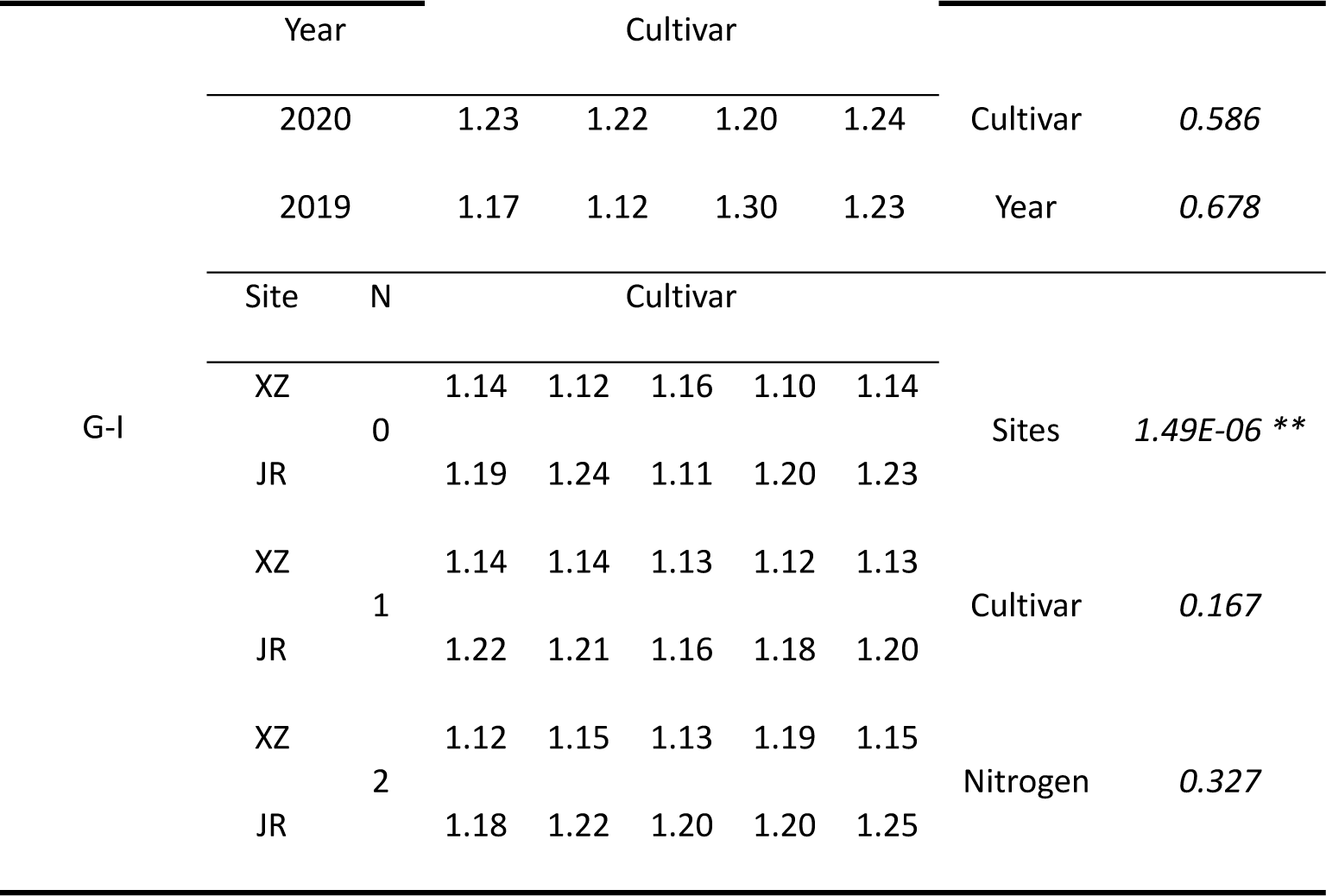
ANOVA analysis results about blade length parameter.

The corresponding between the photothermal quotient and blade length, blade width can be expressed as Y = 0.28 X + 0.83, Y = 0.30 X + 0.79 with R^2^ = 0.49, R^2^ = 0.45, respectively (Fig 2). Therefore, the model based on the growth law of successive phytomers can be described as shown in Table 3, which only needs to input radiation, temperature, the final number of leaves, and the maximum leaf dimensions as model parameters to obtain the detailed dimensions of phytomers.

**Fig. 2.**
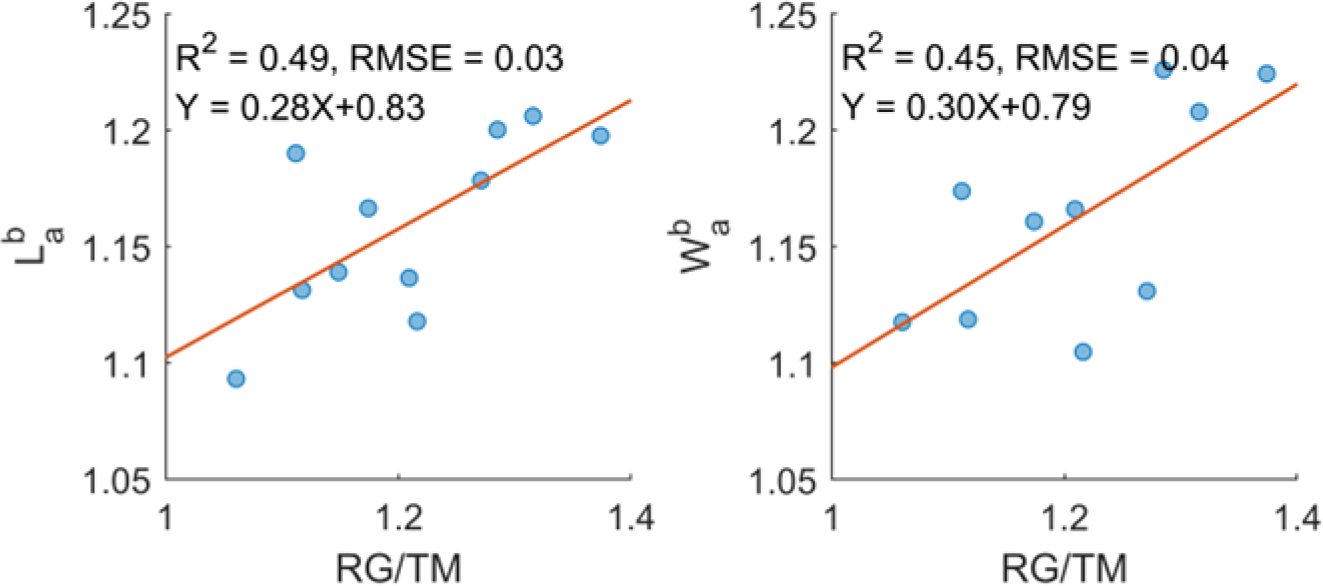
Correlations between blade length 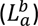, blade width 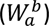 and photothermal quotient.

**Table 3.**
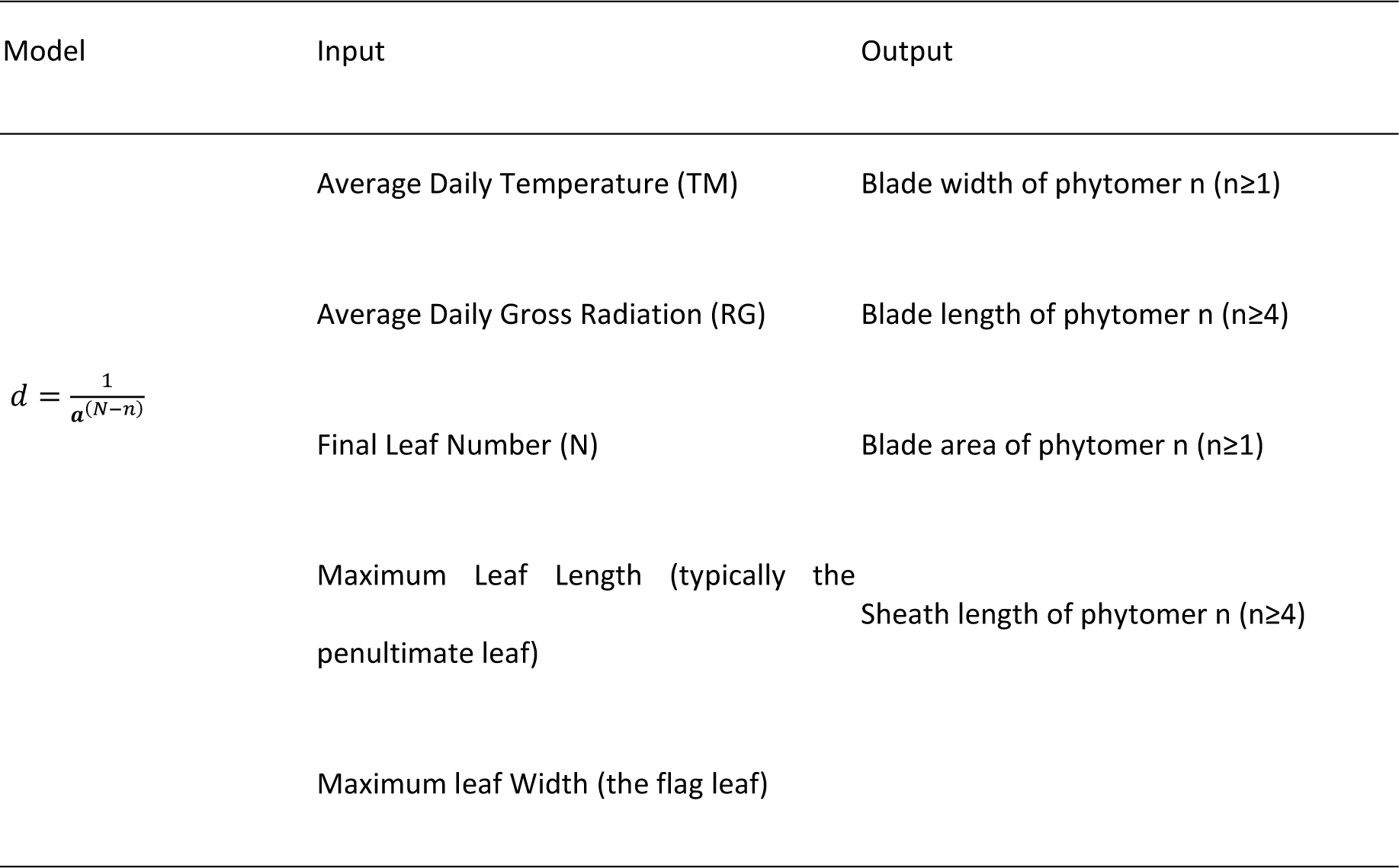
Description of successive pattern model: Input and Output.

| Model | Input | Output |
| --- | --- | --- |
| $d = \frac{1}{a^{(N-n)}}$ | Average Daily Temperature (TM) | Blade width of phytomer n (n≥1) |
|  | Average Daily Gross Radiation (RG) | Blade length of phytomer n (n≥4) |
|  | Final Leaf Number (N) | Blade area of phytomer n (n≥1) |
|  | Maximum Leaf Length (typically the penultimate leaf) | Sheath length of phytomer n (n≥4) |
|  | Maximum leaf Width (the flag leaf) |  |

This study found a strong linear relationship between the final leaf width and the leaf sheath length (Fig 3), which is highly consistent with previous research results (Liu et al. 2019; Zhu et al. 2014), and the parameters of its formula are relatively close. This study incorporates a large number of new datasets and refines the relationship again, so that the final leaf width and leaf sheath length can be estimated from each other by their relationship.

**Fig. 3.**
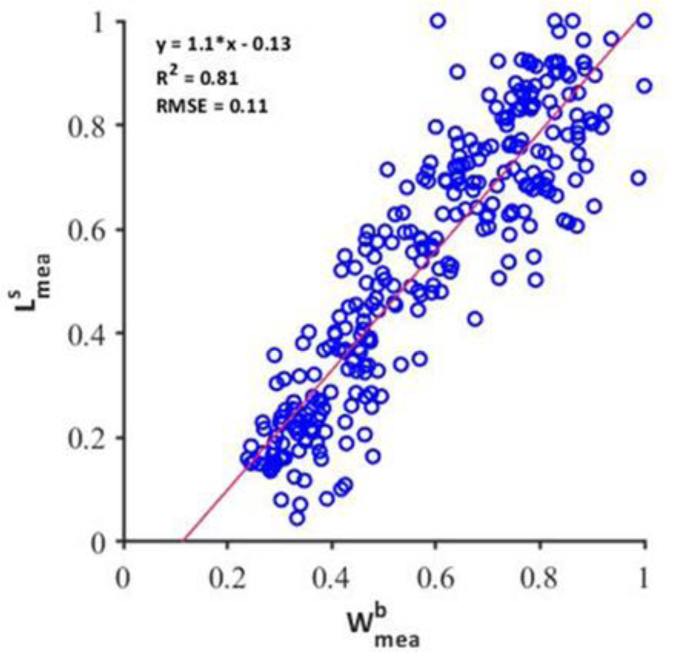
Model performance against final organ dimensions.

### 6.3. Validation of series models

#### 6.3.1. Validation of successive phytomer model for cultivars

According to the results of ANOVA, this study conducted an independent cultivar validation for experiment J (see Table 1). The analysis was modeled with the same 4 cultivars as in experiment I, and the remaining cultivars were validated (cultivars number is more than 90).

Fig. 4 a shows the relationship between the simulated value and the measured value of the blade width, which is uniformly distributed near the red 1:1 line with R^2^ = 0.95 and RMSE = 0.05. Likewise, the model can perfectly predict the sheath length (Fig. 4 b), blade length (Fig. 4 c) and lamina area (Fig. 4 d). This is highly consistent with the ANOVA results that the model’s performance is stable across cultivars.

**Fig 4.**
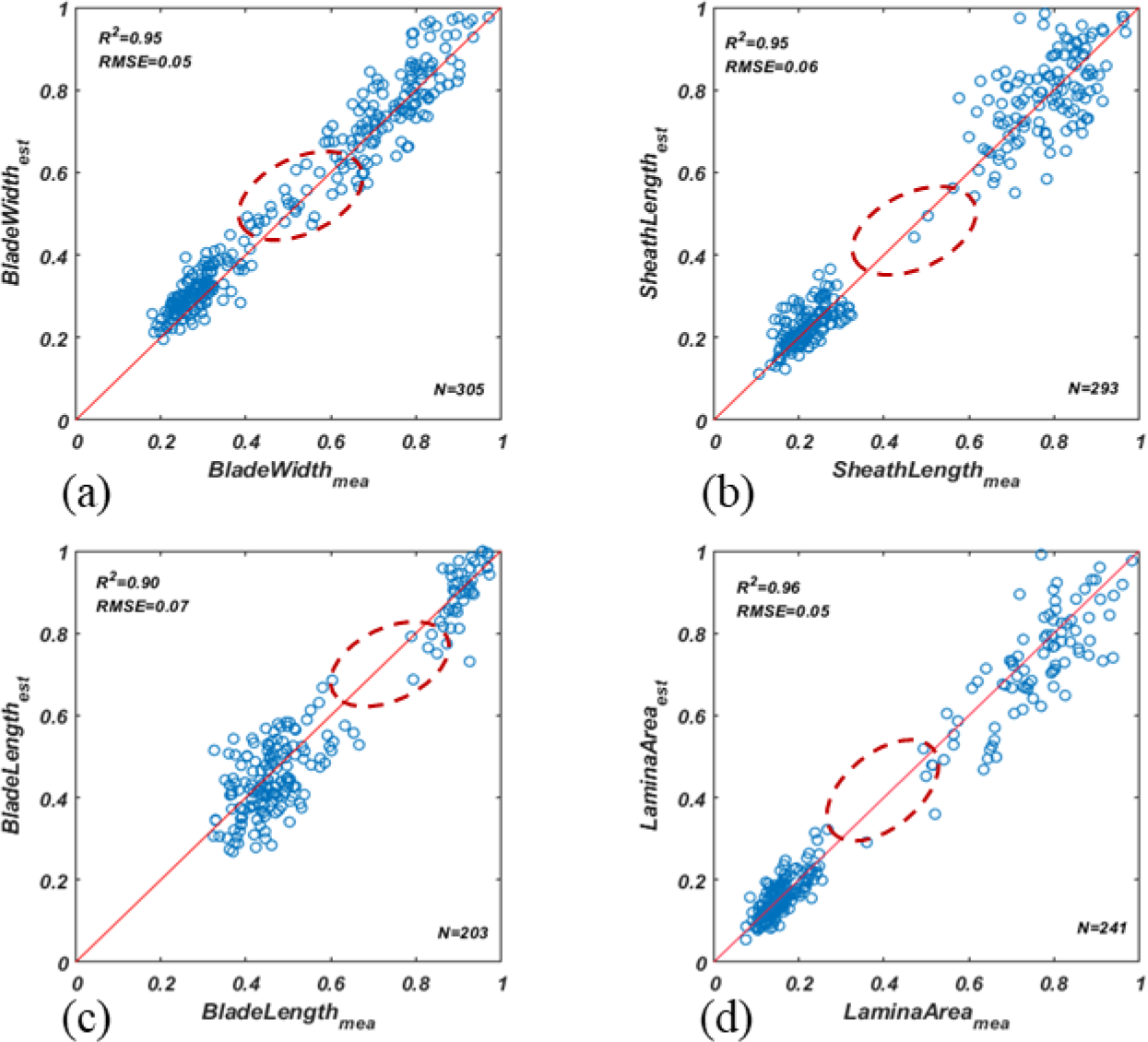
Model performance against final organ dimensions, including blade width (a), sheath length (b), blade length (c), and lamina area (d). The red dot circle shows the missing leaves in 2019 Baima experiment.

#### 6.3.1. Validation of successive phytomer model by sites

We used the hold out method (Hold out 30%) to perform site-by-site validation on the dataset, and the results are shown in Figure 4.

The validation results of the empirical model showed that the simulated and true values of the blade width (Fig 5 a) are uniformly distributed around the 1:1 line. Its accuracy (R^2^ = 0.90, RMSE = 0.06) was higher than the prediction performance for blade length (Fig 5 b) and sheath length (Fig 5 c). Among them, the model’s prediction of sheath length was more scattered on the first several phytomer ranks (Fig 5 c).

**Fig 5.**
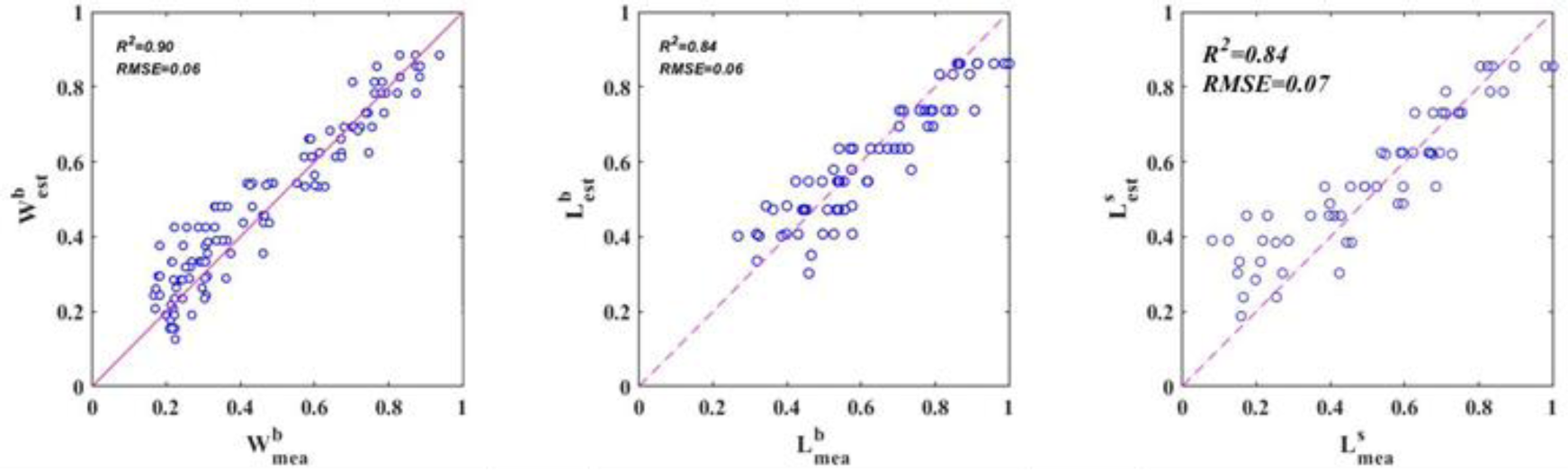
Model performance against final organ dimensions, including blade width, blade length, and sheath length.

#### 6.3.2. Validation of successive phytomer model coupling with meteorological factors

According to the independent experiments A-I, the data sets under more than 10 different meteorology conditions are used for the empirical relationship established in section 6.2 to carry out Leave-One-Out-Cross-Validation (LOOCV), the result is shown in Fig 6.

**Fig.6.**
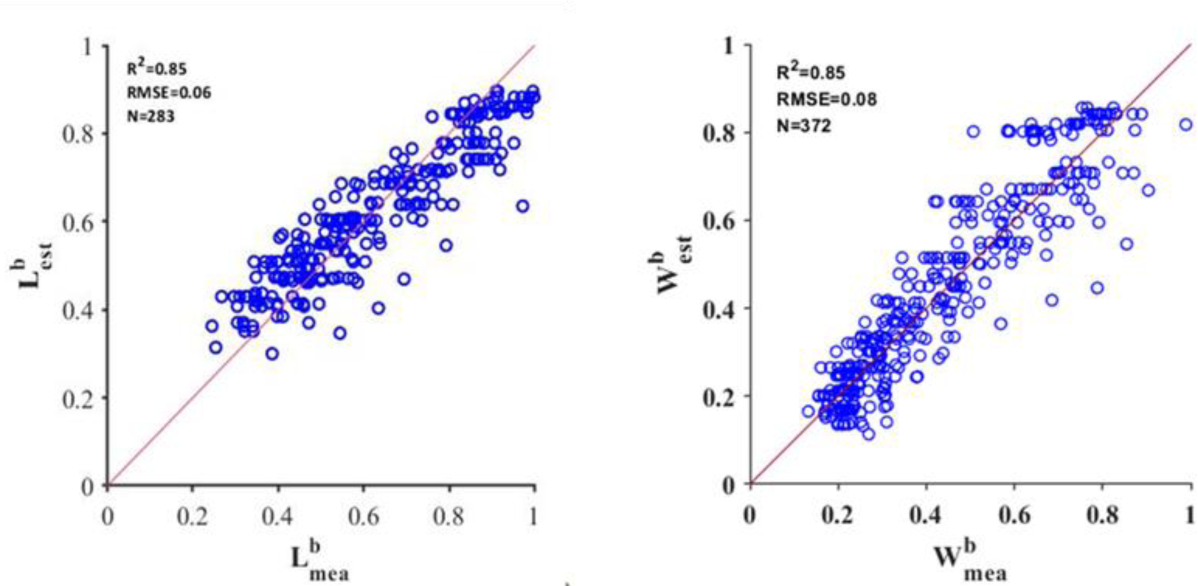
Scatter plots between the estimated value and measured value of blade length (left) / blade width (right). The red line represents the 1:1 fitted line.

The results show that the average daily thermal quotient can more accurately estimate the model parameters and be used for accurate prediction of blade length and blade width, with R^2^ = 0.85.

### 6.4. Comparison with Adel-Wheat and Sirius Quality

#### 6.4.1. Comparison of simulation performance of ADEL-Wheat for blade width

A comparative evaluation of ADEL-Wheat and the two newly proposed models showed that, compared with ADEL-Wheat (Fig 7 A), the model based on the successive growth law significantly improves the prediction accuracy of blade width in all phytomers of the main stem during the entire growth cycle of wheat. R^2^ increased from 0.70 to 0.85, RMSE dropped significantly from 0.2 to 0.08, and all data points were roughly evenly distributed around the 1:1 line (Figure 7 B). The model based on the successive phytomers growth law proposed in this study is more effective for simulating leaf width. The simulation results of ADEL-Wheat for blade width show that although the overall observation results are not significantly overestimated or underestimated, the prediction results of blade width are more convergent in the early stage, and more divergent in the later stage with some outlier observations (Fig. 7 A).

**Fig 7.**
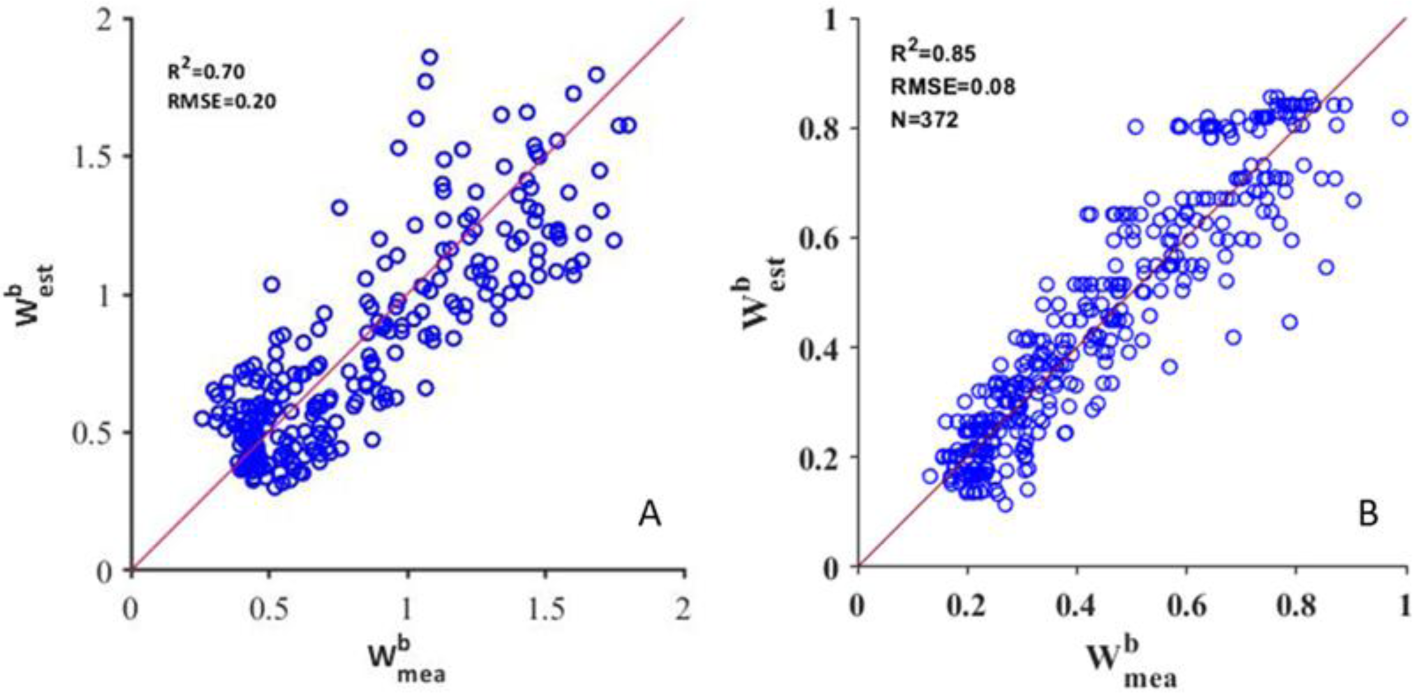
Comparison between observations and estimations of blade width from main stem all phytomer ranks based on ADEL-Wheat (A) and successive law (B) model.

#### 6.4.2. Comparison of Sirius Quality simulation performance for blade area

For early leaves, Sirius Quality determines the constant value of blade area based on empirical parameters of wheat cultivars during modeling, rather than estimating them. The results of this study showed that the blade area of early wheat was significantly different in the skewed distribution (Figure 8).

**Fig 8.**
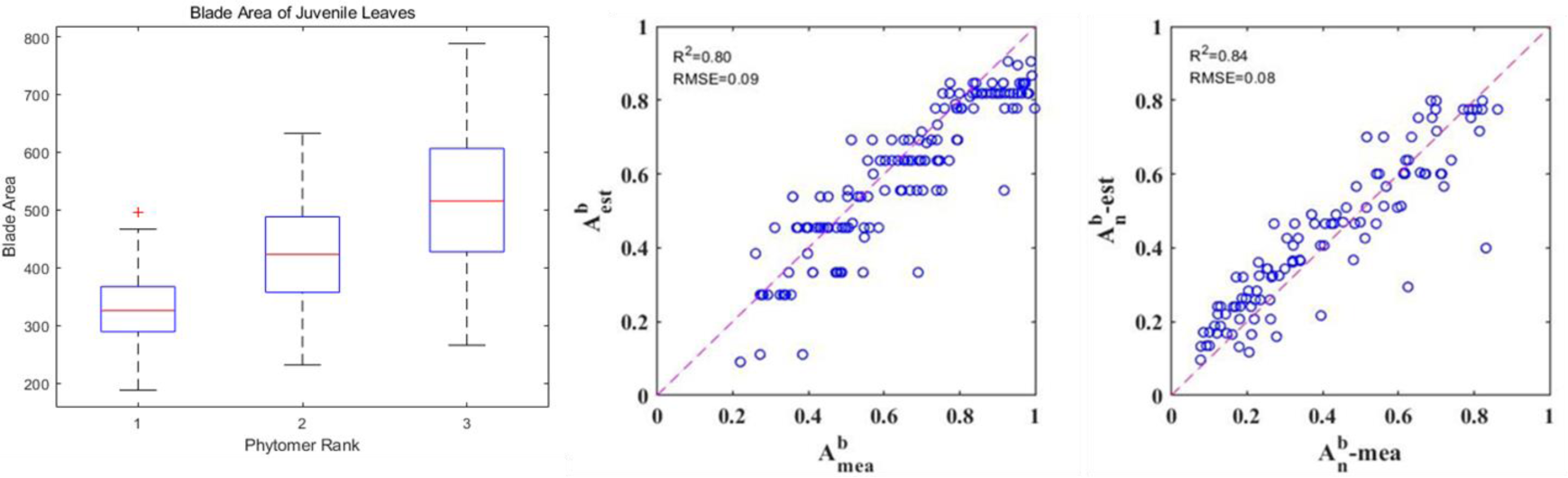
Boxplots of blade area from juvenile phytomers (left), and comparison between observations and estimations of blade width from main stem phytomer ranks based on Sirius Quality (middle) and successive law model (right).

Since Sirius Quality is limited to cultivar parameters, this study improves its calculation method (Eq.3) and validates it with the same dataset to compare with the new model. The results show that RMSE of the blade area validation is basically unchanged (RMSE ≈ 0.08), and R^2^ is slightly higher than Sirius Quality (R^2^ = 0.84), but the prediction results of the new model are more evenly and convergently distributed on the 1:1 line (Figure 8).

## 7. Discussion

### 7.1. The coordination between the dimension of successive organs

According to the work presented by Abbe et al. (1941), we hypothesize that the relationship between the dimension of successive organs determined on the relationship of cell number to a great extent. The same pattern was observed in maize, the blade of which was successively 1.09±0.01 times wider than its predecessor while the circumference of shoot apex is by an average of 1.04±0.02 times. The author directly measured cell numbers and cell size. The result showed that primary condition responsible for the increased size of successive shoot apices is an increase in cell number, while cell size and nuclear size remain essentially uniform. Besides, Gonzalez et al. (2012) reported that cell division and expansion were necessary to form a mature organ. The slopes of cell size and cell number vary with leaves positions on the stem per se, cell number explained largely for the leaf shape since the cell size remains constant relatively (ASHBY and WANGERMANN 1950). This suggests that there is a strong physiological correlation between the two consecutives adjacent phytomers. Besides, Mariem (2016) conclude that previously emerged leaves (length of its whorl) impact the size of the growth zone and mature zone of the leaves that grow within, also the time starts perceiving external stimuli. The relationship was quantified in this study and used for modeling the main stem structure, which improves the authenticity of the structural description of the crop structure function model. The current wheat structure model is mostly based on the phyllochron, driven by thermal time, and is greatly affected by the environment and varieties, and the plant physiological interpretation found in this study has verified the strong robustness of the model. Given the quantified correlation between wheat successive phytomers in this study, other grasses or plants with similar growth patterns have yet to be investigated and quantified, and differences between gramineous species deserve further study.

### 7.2. Environmental and varietal effect on the model performance

#### 7.2.1. The varietal and environmental dependency of the model parameters (complexity and accuracy of the model)

As the ANOVA results showed, the slopes of this pattern did not significantly vary among cultivars under the same treatment. It is probably due to the inherent character of wheat stem as discussed in 7.1 part and the morphology traits that are already shaped by the environment and management they are grow in.

In this study, it is found that model parameters are relevant with the ratio of daily average total radiation to daily average temperature, which is better than the average daily total radiation or temperature effect alone. This results from temperature data used in the analysis partly, which is the air temperature. The length and width of mature leaves depends largely on the size of the vascular bundle that provides nutrients to the leaves. This is based on the previously observed leaf bud primordial base developing upward through the apical meristem long before the new leaf bud primordial group is visible at the apex, and the growth pattern of the leaf is quite stable, so the size of the leaf depends largely on its initial stage (Pieters and Noort 1988). Study has also shown that the occurrence of wheat leaves is mainly dependent on the leaf tip bud temperature (Savvides et al. 2017), even if there is a large difference between the leaf tip temperature and the air temperature plant grows in.

Light radiation itself does not explain the differences in the slope of the successive phytomer sizes. However, studies have shown that the relationship between leaf sheath length and leaf length is related to the emergence system that governs leaf elongation. When the leaf tip emerges from the sheath of a previous phytomer, changes in the environment (especially light interception) affect the final size of the new plant phytomer (Casey et al. 1999). Leaf elongation rate increases linearly with temperature and is not affected by light intensity, while the maximum blade width is not affected by temperature or light intensity. Lacube et al. (2017) has reported the leaf widening was sensitive to whole-plant light intercept, while the leaf elongation was not. Leaf elongation duration is closely related to phyllochron, although this relationship is slightly altered by light intensity (Bos and Neuteboom 1998). In addition, low temperatures and high radiation help plants capture energy (Karafil et al. 2016). This is consistent with the results of this study.

#### 7.2.2. The reality of the model or not properly calibrated

As the profile showed in figure 1, wheat final blade length on main stem reached its peak before the flag leaf, which is typically the penultimate leaf in this dataset corresponding with the description by E.J.M (2002). However, it was reported that the final blade length could decline from the top three to four leaves under various treatments (Evers et al. 2005). The model proposed in this study omitted the relationship between the longest leaf to the flag leaf. Supplementary work focusing on the pattern of blade length towards leaf position after reach the longest one still needs to be done. Besides, the blade and sheath sizes at juvenile stage of wheat deserve to be further described and quantified.

### 7.3. Performance comparison

The blade width derived from ADEL-Wheat is based on the relationship between blade length and blade width, while the model proposed in this study is followed by the spontaneous event. The expanding leaf blades of dicotyledonous species increase both in length and width, whereas a visible leaf of a monocotyledonous species increases only in length, because the width remains unchanged once it has emerged from the sheath tube (Dale 1988). Regarding this, we hypothesize that results in final blade width estimation are better than that of length mainly because that the leaf widening happens in sheath tube, which is affected by plant per se other than external environment. Compared with it, leaf length perceives the external environment once it emerged from sheath then elongates for a relatively long time. The model based on successive pattern provides slightly better blade area estimations than the Sirius Quality, which means the blade area along the phytomer rank is not exactly the linear relationship. According to the calculation format proposed based on the blade sizes pattern, the relationship between blade area and phytomer rank is more exponential than linear.

### 7.4. The minimum set of measurements to characterize the canopy structure and potential for HTP canopy structure

This study introduced a promising model that complements FSPMs and provided insights into high throughput phenotyping canopy structure in more detail. According to the model, only two sets of measurements are enough for characterizing the main stem structure. One happens in the juvenile stage, when to measure the sizes of small leaves. The other one is to measure the uppermost phytomer, more specifically the maximum blade length to the length of the flag leaf and the blade width, internode length of the flag leaf. Since the tillers behave similarly to the main stem, canopy structure can be described by the main stem structure. The upper bigger leaves are much more easily to be identified by the sensors which illustrates that this method is potential for HTP canopy structure. In addition, the method fills the gap in the detailed structure of the wheat canopy observed from the top and can be integrated into the D3P platform (Liu et al. 2019) for in-silico experiments.

## 8. Conclusion

By using the dimension of the final phytomer (and the length of the penultimate leaf) and final phytomer number combined with a coefficient (derived from average daily gross radiation and temperature), the dimension of elements from the phytomer can be estimated by a new model based on well-defined successive patterns (under relatively sufficient nitrogen supply), which is promising for high throughput phenotyping methods, and a clear botanical interpretation from growing apex – cell number rather than size counts largely for the final dimensions of phytomer components.

## 9. Supplementary Data

**Table.S5.** Uncertainty interval of the Eq.4. The variability range of slopes is corresponding to the uncertainty interval, which reveals that *a* value is relatively stable among the experiments in this study. (This may due to the environments of all the experiments are suitable for winter wheat.)

| Slope | Variability |  |  | Uncertainty Interval (95%confidence bounds) |  |  |  |
| --- | --- | --- | --- | --- | --- | --- | --- |
|  | average | range | interquartile range | min | Q1 | Q3 | max |
| $W_b$ | 1.16 | 0.12 | 0.06 | 0.04 | 0.04 | 0.08 | 0.13 |
| $L_b$ | 1.16 | 0.11 | 0.05 | 0.04 | 0.06 | 0.10 | 0.13 |
| $L_s$ | 1.18 | 0.12 | 0.02 | 0.05 | 0.06 | 0.14 | 0.26 |
| $L_i$ | 1.57 | 0.15 | 0.04 | 0.11 | 0.12 | 0.33 | 0.49 |

**Table.S4.**
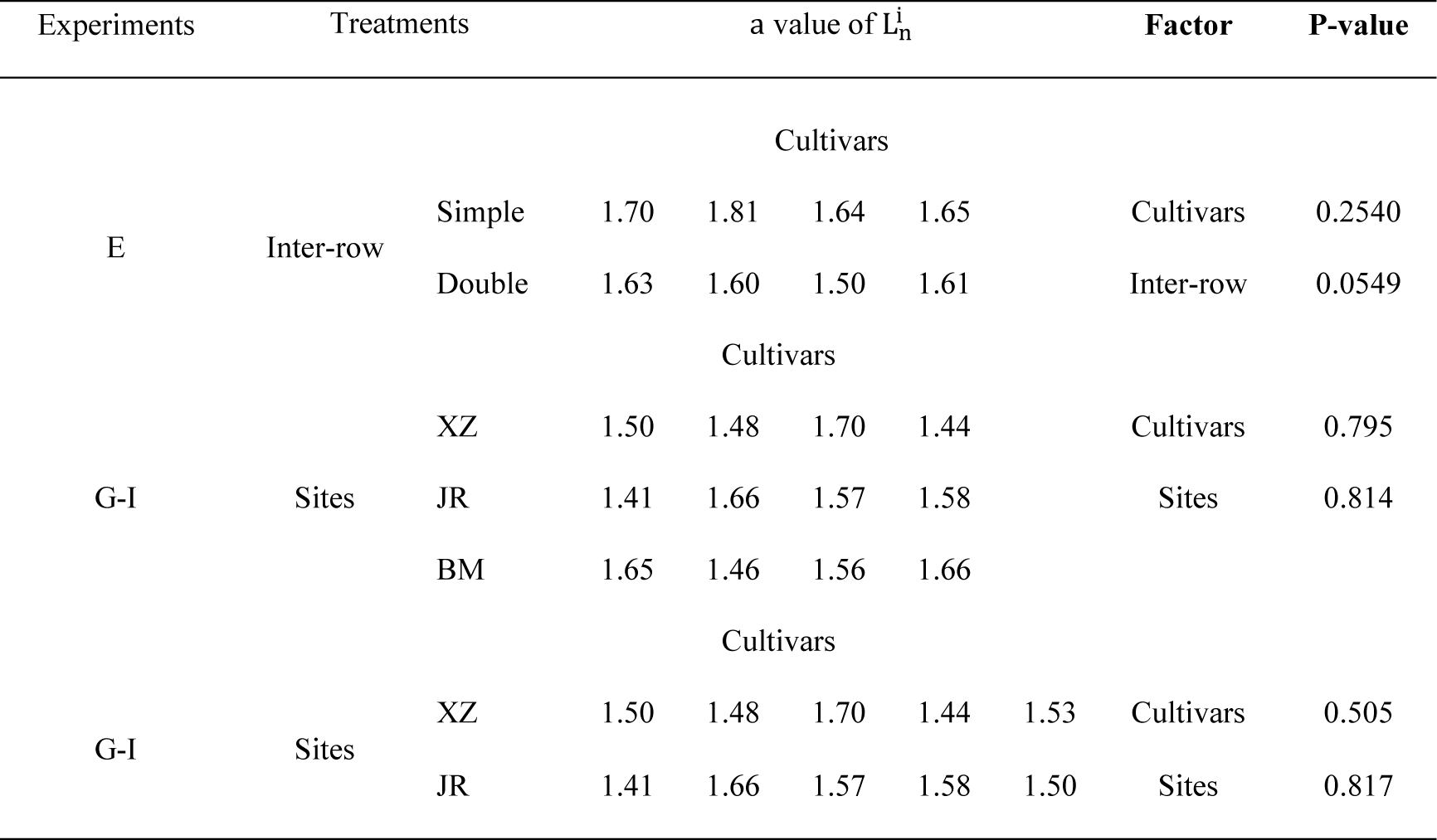
ANOVA analysis results about internode length parameter. Due to the relatively fewer data of internode length and no significant factors were found, there is no detail result about estimation of internode length in this paper

**Table.S3.**
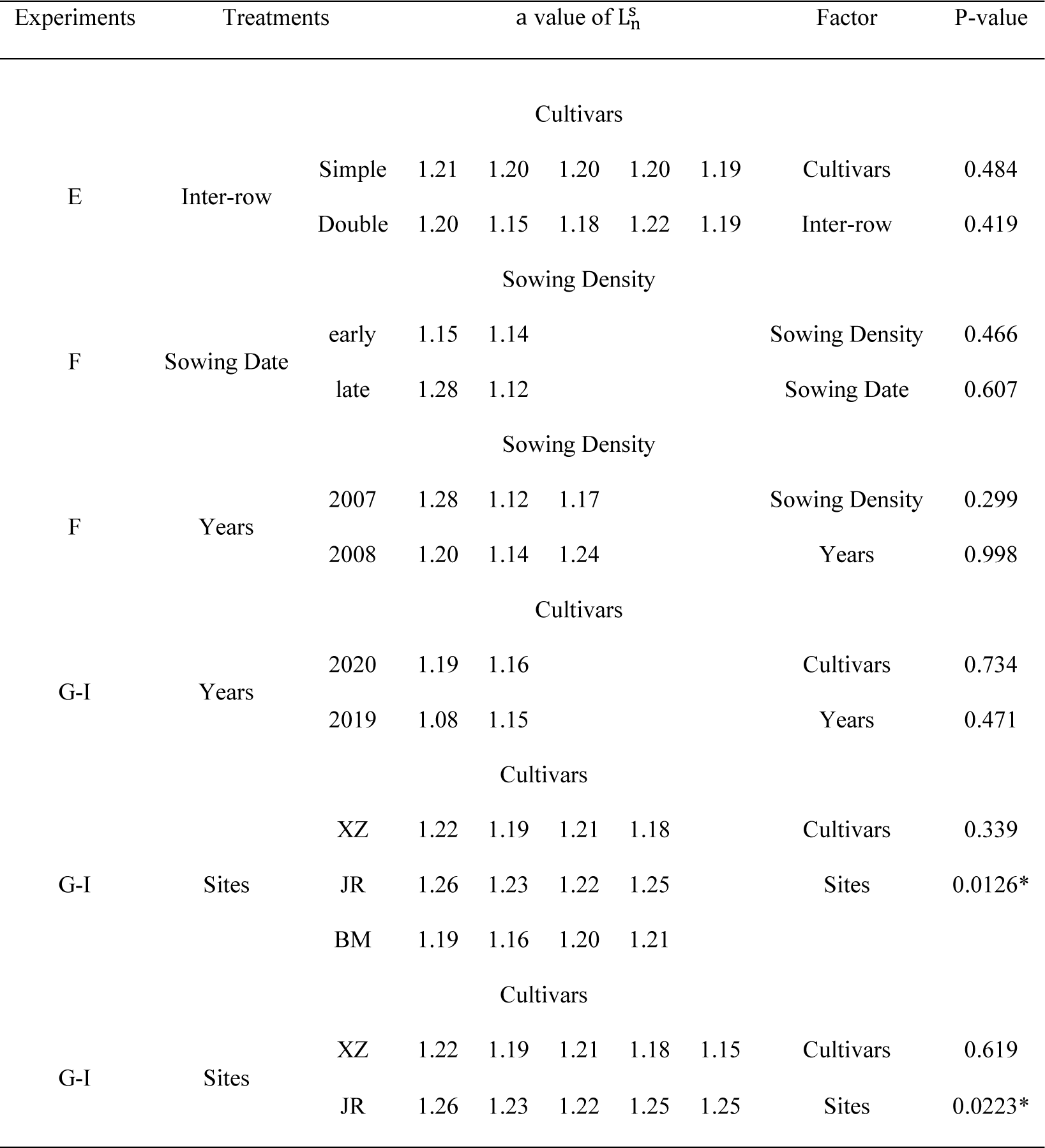
ANOVA analysis results about sheath length parameter.

**Table.S2.**
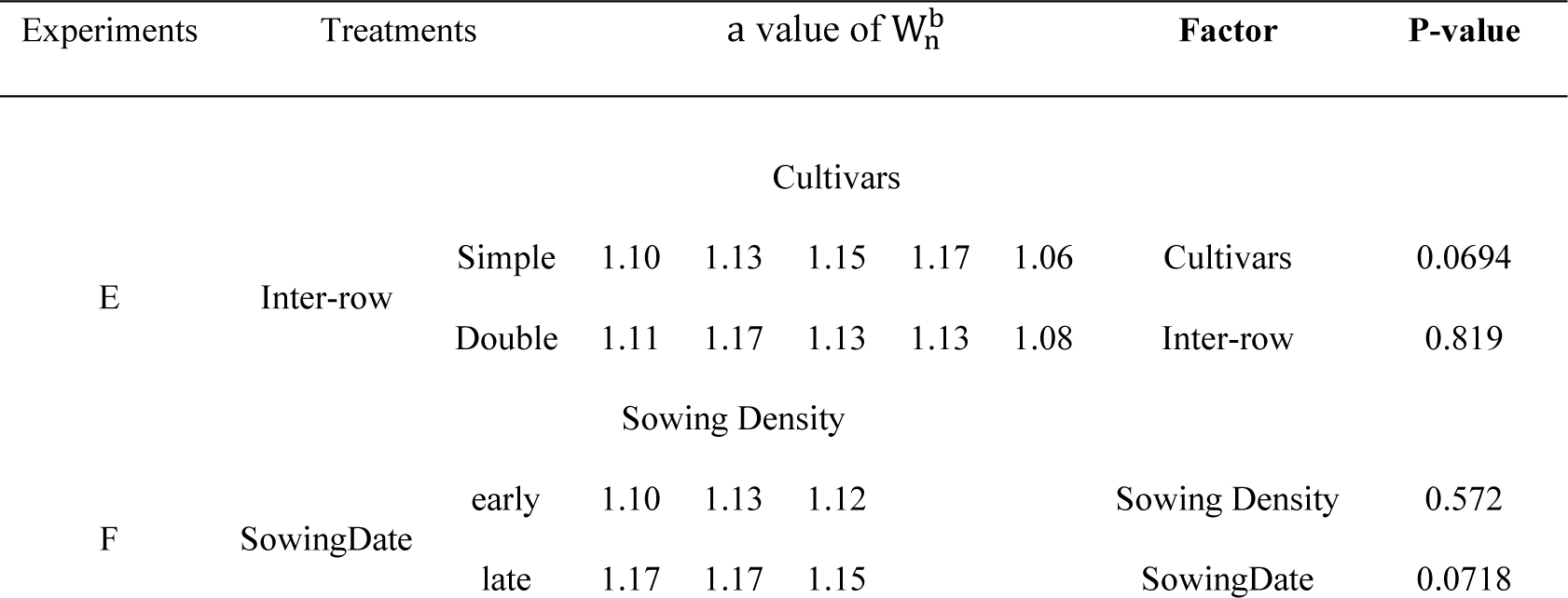
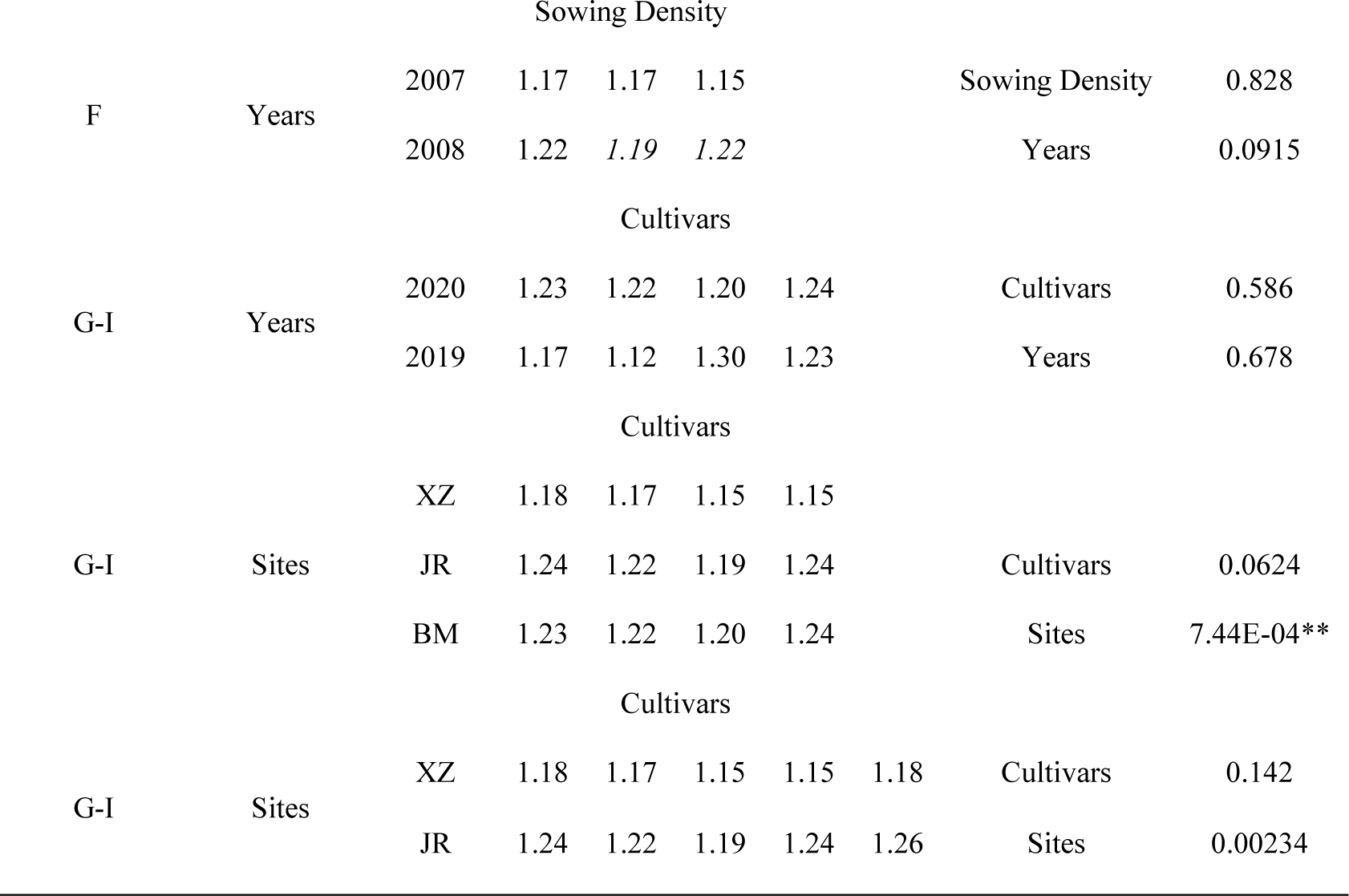
ANOVA analysis results about blade width parameter.

**Fig.S1.**
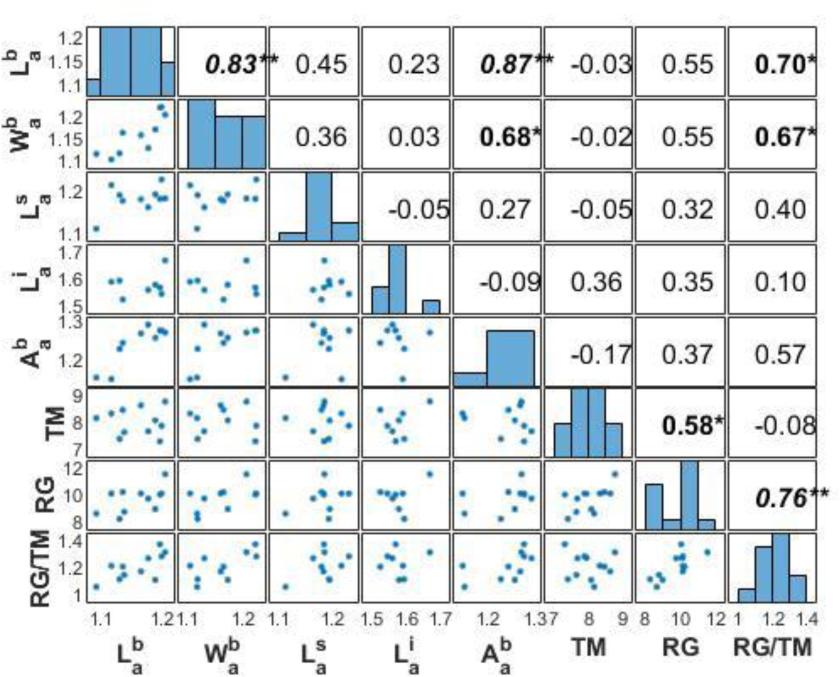
Correlations among organ parameters and meteorological factor.

**Table.S1.**
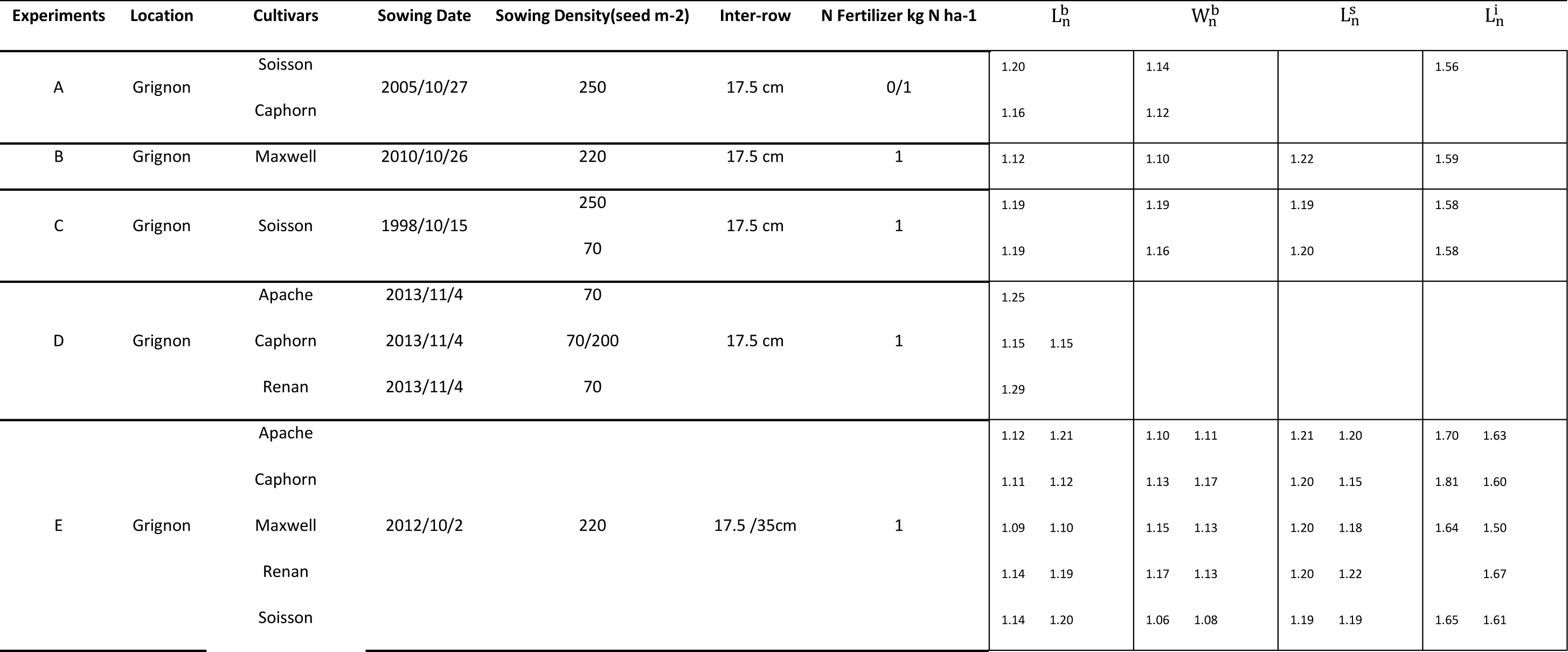
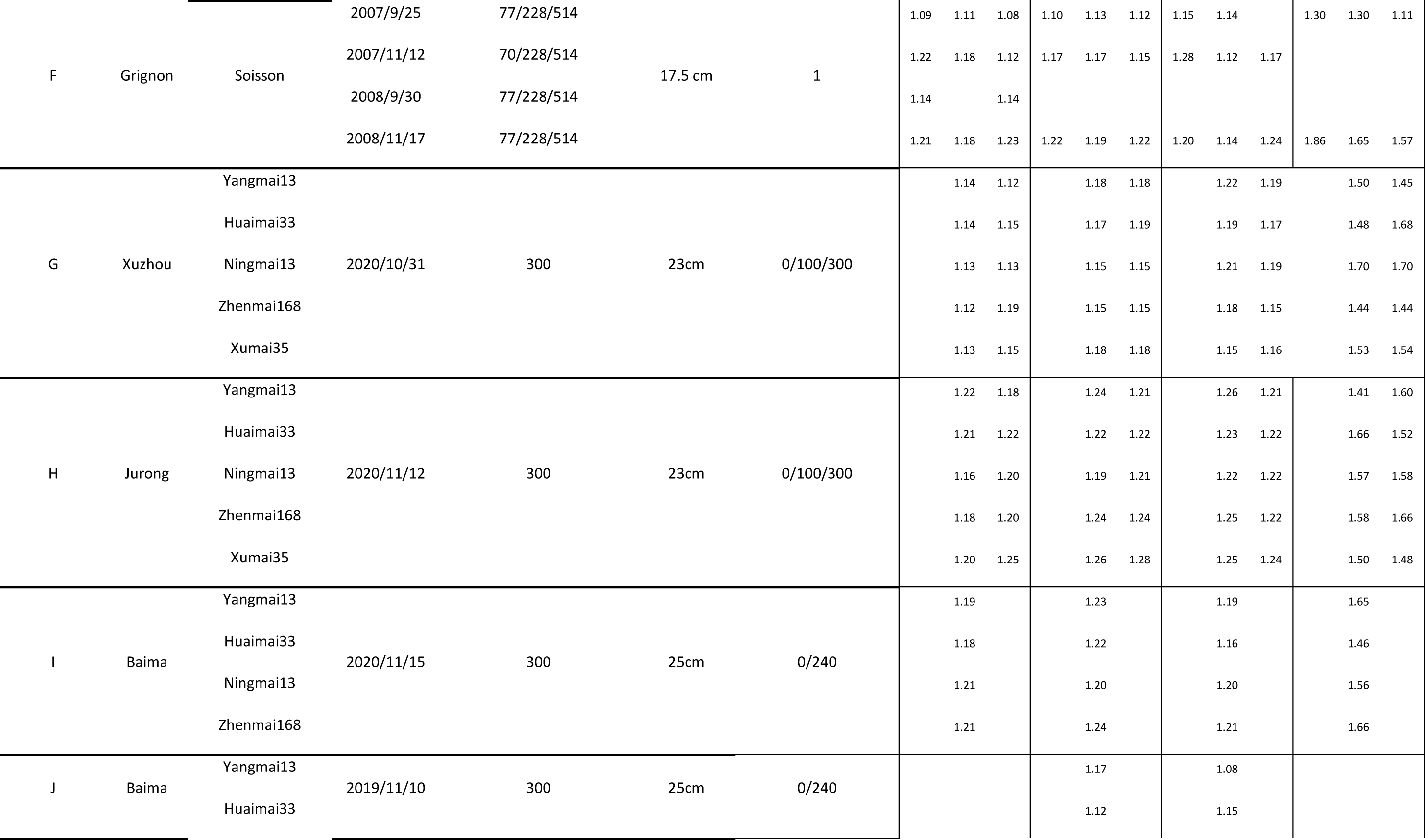
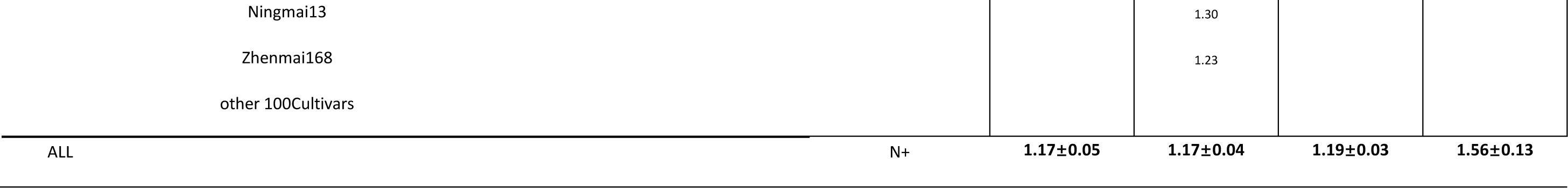
Detailed coefficient a of all treatments.

## 10. Acknowledgments

## Author contributions and acknowledgement

During the entire work, I conducted the field experiment 2019-2021, collected and analyzed the data, then wrote this manuscript. Chen Zhu helped in data collecting and manuscript completing. Qing Li helped in designing Baima field experiments. Pengyan Li, Chaoyu Fan and Yangmingrui Gao helped in data collection.

Thanks to Tiancheng Yang, Ruowen Liu, Jiangmei Mao, Mingxia Dong and Xiaohai Zhan who are involved in data collecting. I am grateful for the data support of Shouyang Liu, Bruno Andrieu from French fields (INRAE) and field experiments support from Dong Jiang and Jing Wang in China. Special thanks to Frederic BARET (INRAE) for his expertise and support for this work.

## 10.1. Conflicts of interest

The author has no conflicts of interest.

## 10.2. Funding

The financial support is from Nanjing Agriculture University.

## 10.3. Data Availability

Dataset and codes are all available if requested.

## Notes

### Competing Interest Statement

The authors have declared no competing interest.

